# Near-infrared imaging of phytochrome-derived autofluorescence in plant nuclei

**DOI:** 10.1101/2023.09.18.558255

**Authors:** Akira Yoshinari, Reika Isoda, Noriyoshi Yagi, Yoshikatsu Sato, Jelmer J. Lindeboom, David W. Ehrhardt, Wolf B. Frommer, Masayoshi Nakamura

## Abstract

Capturing images of the nuclear dynamics within live cells is an essential technique for comprehending the intricate biological processes inherent to plant cell nuclei. While various methods exist for imaging nuclei, including combining fluorescent proteins and dyes with microscopy, there is a dearth of commercially available dyes for live-cell imaging. In *Arabidopsis thaliana*, we discovered that nuclei emit autofluorescence in the near-infrared (NIR) range of the spectrum and devised a non-invasive technique for the visualization of live cell nuclei using this inherent NIR autofluorescence. Our studies demonstrated the capability of the NIR imaging technique to visualize the dynamic behavior of nuclei within primary roots, root hairs, and pollen tubes, which are tissues that harbor a limited number of other organelles displaying autofluorescence. We further demonstrated the applicability of NIR autofluorescence imaging in various other tissues by incorporating fluorescence lifetime imaging techniques. Nuclear autofluorescence was also detected across a wide range of plant species, enabling analyses without the need for transformation. The nuclear autofluorescence in the NIR wavelength range was not observed in animal or yeast cells. Genetic analysis revealed that this autofluorescence was caused by the phytochrome protein. Our studies demonstrated that nuclear autofluorescence imaging can be effectively employed not only in model plants but also for studying nuclei in non-model plant species.

## Introduction

The nucleus is a cell organelle that plays a pivotal role in many biological processes, including gene expression, DNA replication, and cell division (Dundr and Misteli, 2001), that are crucial for plant growth and development. Eukaryotic cells exhibit a significant morphological plasticity and dynamic migration of the nucleus. In plants, morphology and positioning of the nucleus vary depending on tissue types (Chytilova et al. 2000; Pronina et al. 2022). To understand the structure and function of this organelle, nuclear visualization techniques have been developed. Nuclei have been extensively stained using classical dyes such as 4’,6-diamidino-2-phenylindole (DAPI) and Hoechst33342 (Kapuscinski, 1995; Latt and Wohlleb, 1975). These dyes are not fully functional in unfixed tissues and are therefore predominantly used to stain fixed cells that are then imaged using fluorescence microscopy and confocal laser microscopy. Although DNA dyes, including the SYTO^®^ dye series, have also been utilized for nuclear imaging, their cytotoxicity and impermeability to cell membranes make them challenging to use for live cell imaging (Yagi et al. 2021; Yu et al. 1997). Recently, an N-Aryl Pyrido Cyanine derivatives has been developed, and this dye is the only one successfully used for live cell nuclear imaging of Arabidopsis leaves and roots without adverse effects (Uno et al. 2021). With the increasing ease of genetic manipulation in Arabidopsis, numerous fluorescent proteins (FPs) have been fused to nuclear localization signal peptides or nuclear-localized proteins for successful nuclear imaging in living cells. However, these FP-based methods require genetic modification, rendering them impractical for plant species lacking well-established genetic modification protocols. Therefore, there is an urgent need to develop a simple nuclear imaging technique in the field of plant science.

In this study, we developed a nuclear imaging method that uses inherent nuclear autofluorescence. Spectrum scanning analysis revealed that Arabidopsis nuclei have excitation and emission wavelengths in the near-infrared (NIR) range, and that this autofluorescence were easily visualized using conventional confocal laser microscopy. We demonstrated that NIR nuclear autofluorescence allows multi-color live-cell imaging when used simultaneously with the commonly used FPs, Green Fluorescent Protein (GFP) and Red Fluorescent Protein (RFP). Furthermore, this method was used for plant species other than model plants to enable imaging of nuclear dynamics in plant species where genetic modification protocols have not yet been established. NIR nuclear autofluorescence seems to be a plant-specific feature, as it has not been confirmed in animal or yeast cells. We have shown that this autofluorescence was caused by the red/far-red light receptor phytochrome. Nuclear autofluorescence provides a new and simple imaging technique for nuclear dynamics in plant science research and also a new perspective on phytochrome biology.

## Results

### Unexpected autofluorescence of nuclei in *Arabidopsis thaliana*

The root cells of Arabidopsis exhibit diminished autofluorescence relative to cells in the shoot due to lower concentrations of chlorophyll and less lignification in secondary cell walls. Therefore, using root tissues for live-cell fluorescence imaging and implementation of new techniques such as light sheet microscope and chemical labeling tend to be less challenging than in shoot tissues (Iwatate et al. 2020; von Wangenheim et al. 2017). We have found that when roots are excited at a wavelength of 640 nm, autofluorescence can be detected in the near-infrared (NIR) wavelength range of 656–700 nm (Fig. 1A). The fluorescent compartments appeared primarily as round-shaped structures in every cell type (Fig. 1A).

**Figure 1.**
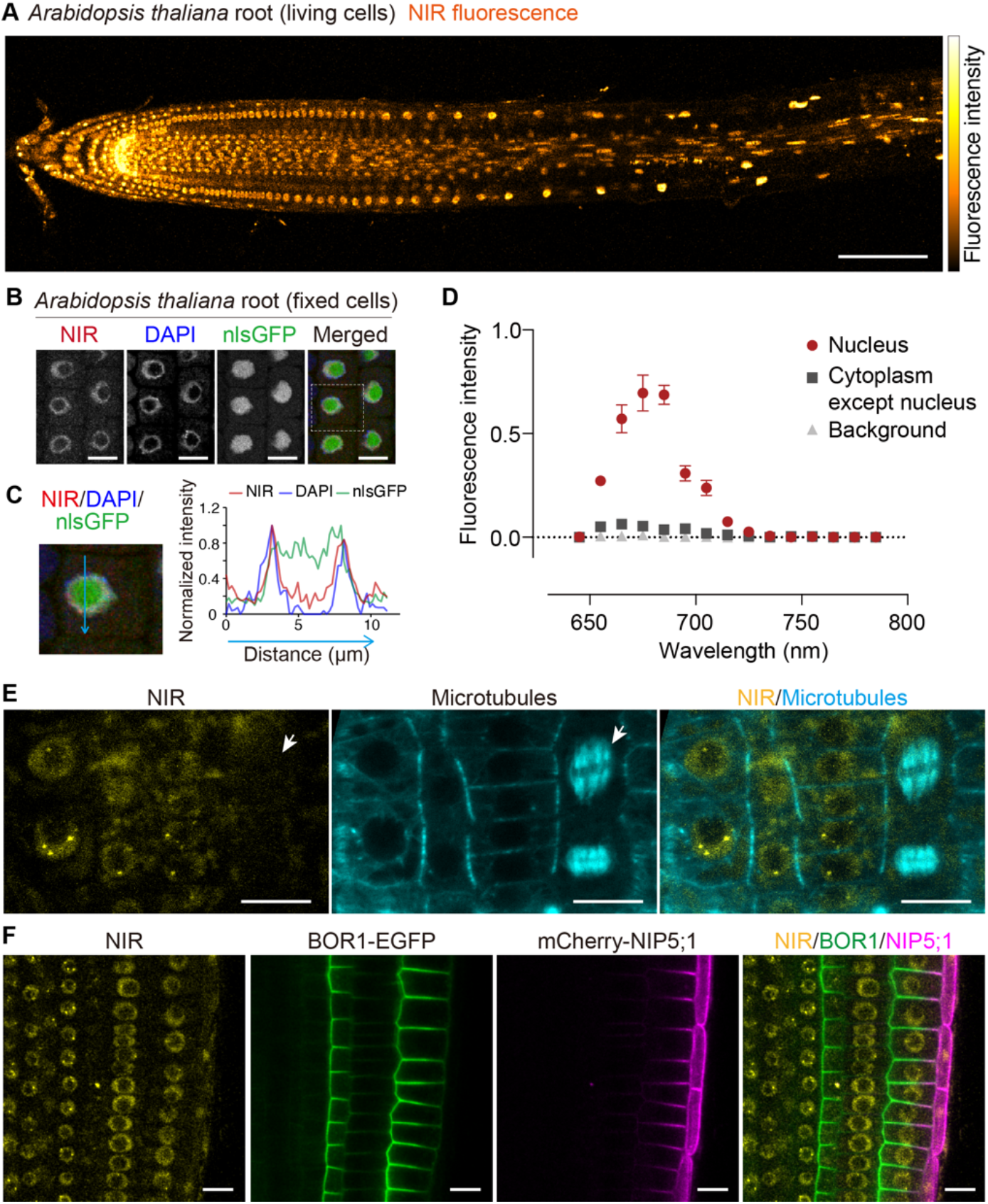
Near-infrared autofluorescence in nucleus. (A) Confocal image of *Arabidopsis thaliana* seedling root was exposed to a laser at the 640 nm wavelength and the fluorescence was detected in the wavelength range of 656–700 nm. The plant was grown on modified MGRL medium containing 1 µM boric acid for 4 days. (B) Fixed Arabidopsis plant expressing nlsGFP, GFP fused with nuclear localization signal, and stained with DAPI. (C) Enlarged image of the indicated area in B. Intensity of each fluorescence along the arrow in the enlarged image was plotted. (D) Fluorescence intensity scan according to different detection ranges. Arabidopsis root was exposed to a laser at 640 nm and the fluorescence was scanned with 10-nm detection window from 645 nm to 795 nm without gap (15 steps). Error bars represent mean ± SE. *n* = 6 (nucleus), 3 (cytoplasm except nucleus), and 3 (background) in one representative root. (E) Multicolor fluorescence imaging of NIR autofluorescence and microtubules (GFP-TUA) in primary root of transgenic Arabidopsis plant grown on half-strength of MS medium for 4 days. Arrows indicate dividing cell where NIR fluorescence was not observed. (F) Multicolor fluorescence imaging of NIR autofluorescence, BOR1-EGFP, and mCherry-NIP5;1 in primary root of transgenic Arabidopsis plant grown on modified MGRL medium containing 0.5 µM boric acid for 4 days. Scale bars represent 100 µm (A), 10 µm (E and F), respectively.

To elucidate the identity of the round-shaped compartments found in each cell, we co-stained with nuclear markers. By using 4’,6-diamidino-2-phenylindole (DAPI) and a nuclear-targeted green fluorescent protein (nlsGFP) in fixed Arabidopsis root cells, we demonstrated that the NIR autofluorescence was derived from nuclear materials (Fig. 1B). Additionally, this autofluorescence was more prominent in the nucleoplasm stained with DAPI than in the nucleolus (Fig. 1C). Within each cell, the NIR autofluorescence detected within the range of 650–720 nm primarily arose from the nuclei (Fig. 1D).

The observance of this nuclear NIR autofluorescence prompted us to explore its potential for multi-color imaging with other fluorescent proteins of different wavelengths. Here, we present examples using NIR autofluorescence in combination with GFP fused to α-tubulin 5 (TUA5) (Fig. 1E), as well as with the enhanced eGFP fused to the borate exporter BOR1 and mCherry fused to the boric acid channel NIP5;1 (Fig. 1F). Interestingly, the intensity of NIR autofluorescence was observed to decrease in mitotic cells compared to interphase cells (Fig. 1E). This was also observed when the NIR autofluorescence was detected at the same time as the nuclear protein markers, histone H2B-mClover (Supplementary Fig. S1). It is worth noting that the expression of *BOR1- EGFP* or *mCherry-NIP5;1* did not significantly interfere with the nuclear NIR fluorescence and that their fluorescence were detected separately without significant optical crosstalk (Fig. 1F). These results demonstrate the feasibility of using nuclear NIR autofluorescence for multicolor imaging in living plant cells.

We also occasionally observed smaller but denser granular structures in the quiescent center, pericycle initials, vasculature initials, and ground tissues with excitation 640 nm wavelength (Supplementary Fig. S2). Similar granular structures were also observed when excited at a violet wavelength (405 nm), and these overlapped with the granules with NIR fluorescence (Supplementary Fig. S2). Since the 405-nm laser did not excite nuclei, the fluorescent molecule observed in the granular structures in the root tip region was likely distinct from that in the nuclei. It should be noted that, in older plants grown over a week, root tissues tended to emit an intense autofluorescence within the cortex (Supplementary Fig. S3). Similar to the granular structures in the root tip region, the autofluorescence in the cortex was detected by excitation with a 405-nm laser (Supplementary Fig. S3), indicating that the cortex-specific autofluorescence observed in older roots was also distinct from the nucleus-specific NIR autofluorescence.

### Nuclear autofluorescence imaging in Arabidopsis

The demonstration of nuclear autofluorescence led us to assess its feasibility for further live-cell imaging of plant cells. We investigated nuclear dynamics in other tissue types, especially pollen and elongating pollen tubes, which, similar to the seedling roots, lack chloroplasts containing the prominent autofluorescent molecule chlorophyll. To evaluate nuclear NIR autofluorescence in pollen, we co-stained DNA with green fluorescent dyes such as Hoechst3342 or SYBR Green I. Our results showed that the NIR autofluorescence was more pronounced in the vegetative nucleus (white arrows in Fig. 2A and B) than in the sperm cell nuclei (gray arrows), whereas the DNA dyes enabled the visualization of one vegetative nucleus, and two small nuclei in the sperm cells (Figs. 2A and 2B). Notably, the NIR autofluorescence facilitated clearer imaging of the vegetative nucleus than the DNA dyes, even during pollen tube elongation (Fig. 2C). We observed the nucleus moving as the pollen tube elongated (Video S1). While it is recommended to use SYBR Green to visualize both vegetative and sperm cell nuclei, the NIR autofluorescence method allowed us to observe the dynamics of the vegetative nucleus in pollen tubes without the aid of dyes. Our observations indicated that the NIR-fluorescent molecules were differentially distributed between the vegetative and sperm nuclei.

**Figure 2.**
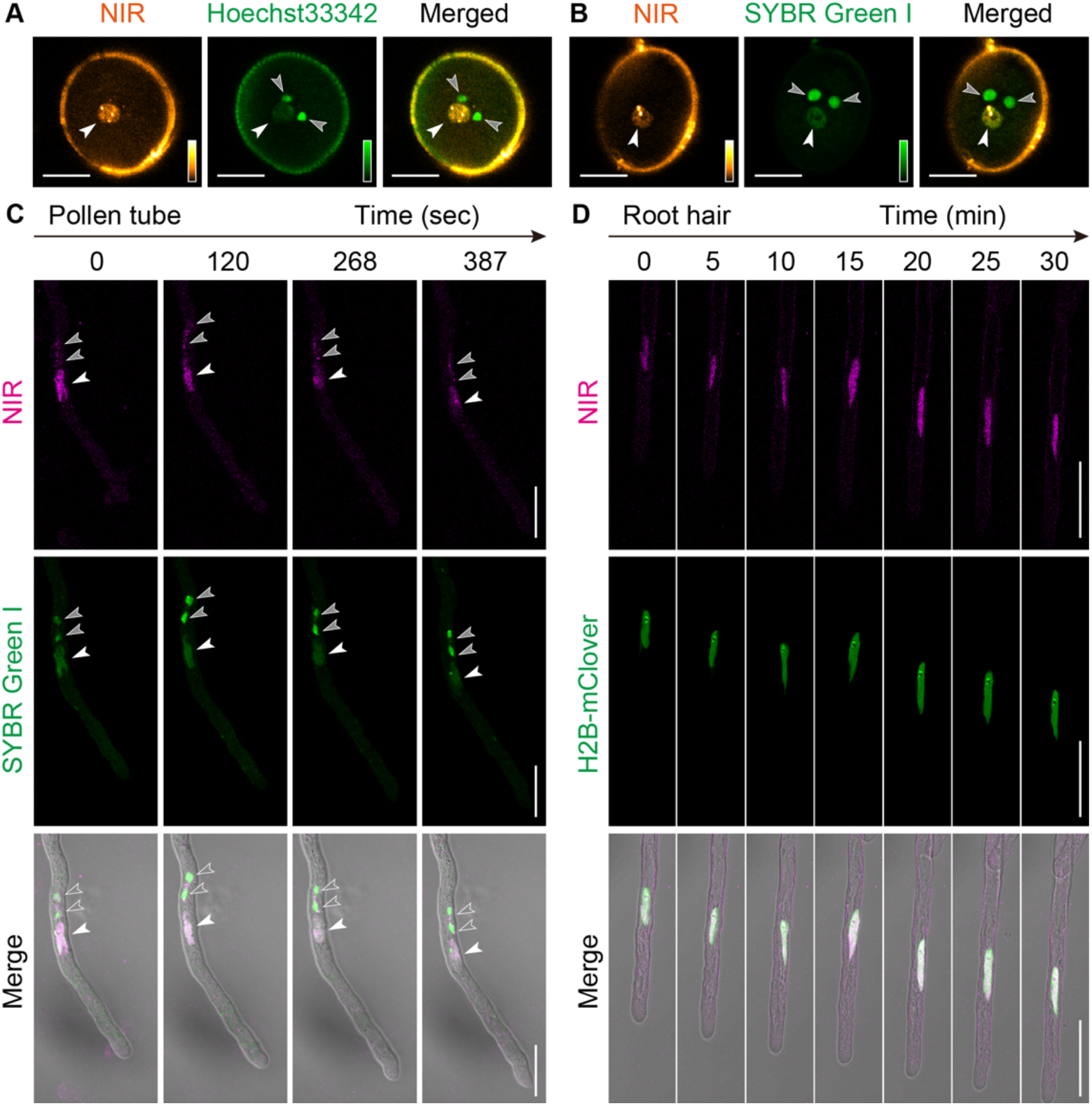
Application of nuclear autofluorescence to live-cell imaging of root hair and pollen tube. (A, B) Near-infrared autofluorescence excited by 640-nm laser and DNA stained by Hoechst33342 or SYBR Green I was excited by 488-nm laser in pollen grain. Gray and white arrowheads indicate nuclei of sperm cells and vegetative cells, respectively. (C) NIR autofluorescence and SYBR Green I observed in pollen tube of wild-type Col-0 plant. The merged images are compound micrographs of bright field transmission and the corresponding NIR and SYBR Green I fluorescence images. (D) NIR autofluorescence observed in root hair of transgenic Arabidopsis plant expressing H2B-mClover grown on modified MGRL medium containing 30 µM boric acid for 4 days. The merged images are compound micrographs of bright field transmission and the corresponding NIR and mClover fluorescence images. Scale bars represent 10 µm (A–C) and 50 µm (D), respectively.

The nuclear autofluorescence imaging technique is also useful tool for monitoring the movement of cell nuclei in root hair. This technique was used to observe the migration of nuclei towards the apical or basal side in Arabidopsis root hair cells (Fig. 2D; Video S2). It is established that the regulation of root hair growth involves maintaining a specific distance between the nucleus and the root hair tip through myosin-mediated mechanisms (Ketelaar et al. 2002). Moreover, it has been reported that myosin-mediated nuclear migration is influenced by light and dark stimuli (Tamura et al. 2013), although the precise molecular mechanism and associated details remain unclear. One notable advantage of this simple autofluorescence imaging approach is that it obviates the need for a fluorescent protein reporter, thereby reducing the time required for introducing a nuclear observation marker into mutant plants. This technique, in turn, should facilitate progress in the field of root hair growth research.

### Nuclear autofluorescence imaging in other plant species

To evaluate the potential of NIR nuclear autofluorescence in other plant species, we utilized confocal microscopy imaging to examine plant species that we routinely use, such as tobacco (*Nicotiana benthamiana*), rice (*Oryza sativa* subsp. *japonica* cv. Nipponbare), onion (*Allium cepa*), and a cultured cell line of *Nicotiana tabacum* BY-2 cells (Fig. 3A). NIR nuclear autofluorescence imaging was readily applicable in these experimental model plants. Next, we collected and imaged seedlings of wild plants, including *Capsella bursa-pastoris* and *Cardamine hirsute*, and crop plants germinated from seeds purchased from a grocery store, including roquette (*Eruca vesicaria* subsp. *sativa*), carrot (*Daucus carota* subsp. *sativus*), coriander (*Coriandrum sativum*), cucumber (*Cucumis sativus*), green onion (*Allium fistulosum*), and spinach (*Spinacia oleracea*) (Fig. 3B). Our results showed that most of these plant species exhibited typical nuclear NIR autofluorescence (Fig 3B), with exceptions for cucumber and spinach seedlings as the seedling roots of *Cucumis sativus* displayed prominent autofluorescence derived from plastids. In spinach, the root cells did not exhibit significant nuclear NIR autofluorescence (Fig. 3B). Instead, strong punctate autofluorescence was observed around the nucleus, thought to emanate from granular organelles presumed to be plastids or mitochondria surrounding the nucleus. This punctate autofluorescence may have interfered with the imaging of nuclear autofluorescence. Together, these observations indicated that nuclear NIR autofluorescence is a common feature in plant species.

**Figure 3.**
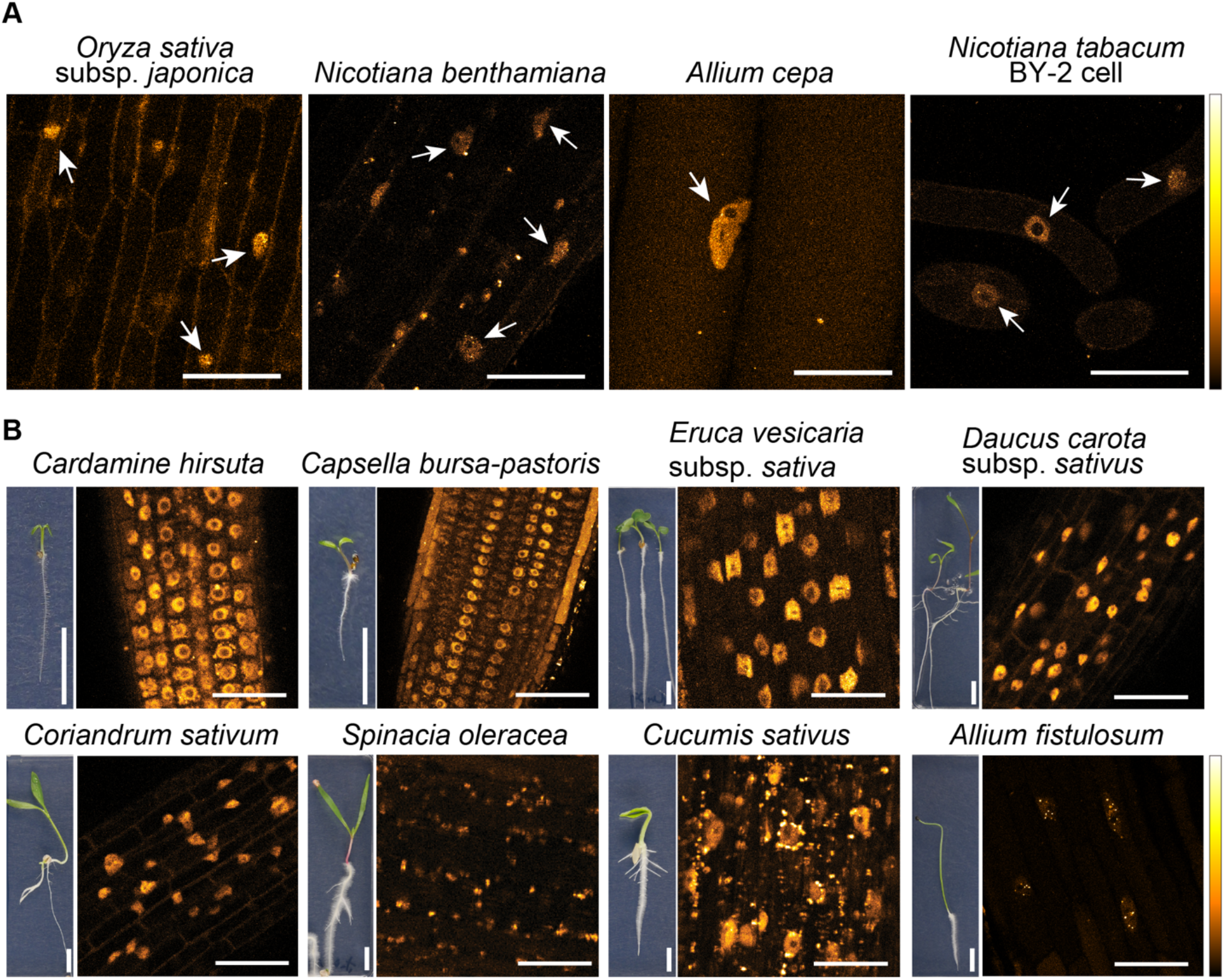
Application of nuclear autofluorescence to live-cell imaging of various plant species. (A) NIR autofluorescence excited by 640-nm laser in model plants including rice (*Oryza sativa* subsp. *Japonica* cv. *Nipponbare*), tobacco (*Nicotiana benthamiana*), onion (*Allium cepa*), cultured cell (tobacco BY-2 cells). Arrows indicate nuclei. (B) Photographs of seedling and NIR autofluorescence images in wild and domesticated non-model plants. Scale bars represent 50 µm (confocal images in A, B) and 10 mm (photographs in B), respectively.

Using the nuclear autofluorescence, we conducted time-lapse fluorescence imaging of nuclei in the roots of non-model plants. The nuclear NIR autofluorescence was observed over 4 hours in growing root of *Cardamine hirsute* (Video S3). Notably, NIR-images with relatively higher signal-to-noise ratios were acquired in roquette roots among the non-model plants (Figs. 4A–C). This outcome can be attributed to their lower levels of background autofluorescence in the other species. The nuclei were observed to transform and migrate within both root tip cells (Fig. 4C; Video S4) and root hair cells (Video S5). In summary, our findings demonstrated that, in many plant species, nuclear NIR autofluorescence is relatively stable and is durable enough to be used for time-lapse imaging in cases where the autofluorescence derived from plastids is not prominent.

**Figure 4.**
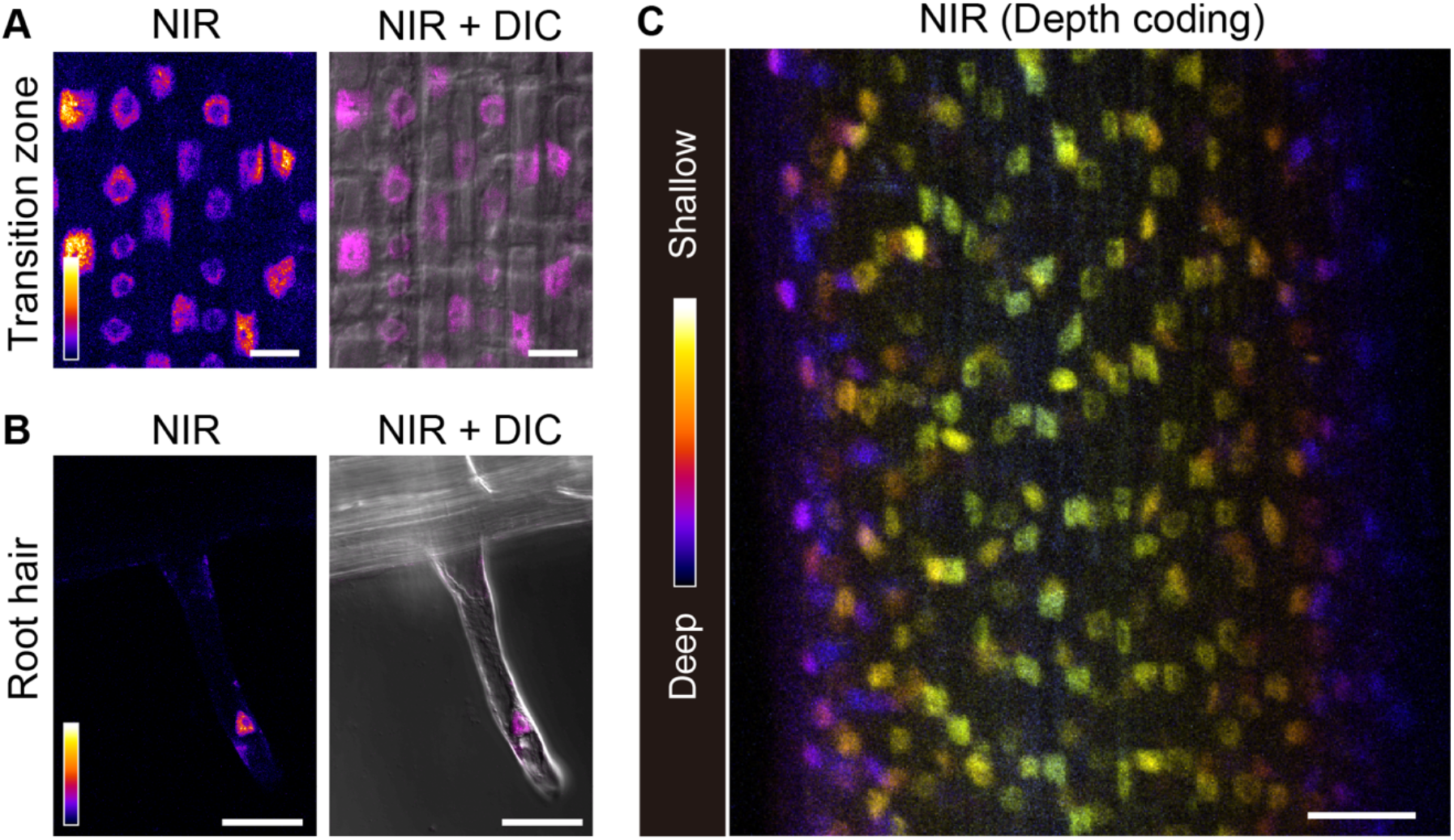
Live-cell imaging of nucleus in roquette root. (A, B) NIR autofluorescence excited by 640-nm laser in root cells and root hair cells in roquette (*Eruca vesicaria* subsp. *sativa*) seedling. (C) A depth-coding image of nuclei based on z-stack images of roquette root. Scale bars represent 20 µm (A) and 50 µm (B, C), respectively.

### Fluorescence lifetime imaging microscopy enables distinction of nuclei from other autofluorescent structures

To further elucidate the characteristics of nuclear autofluorescence, we performed fluorescence-lifetime imaging microscopy (FLIM) in the meristematic epidermal cells of *Arabidopsis thaliana* roots (Fig. 5A). The autofluorescence within nuclei showed little variability in the lifetime of the signal, at 0.67 ± 0.008 ns (Figs. 5B and 5C). Also through the use of FLIM, we successfully identified fluorescence lifetime signals from nuclei embedded in chloroplasts with stronger autofluorescence, as in the moss *Physcomitrium patens* (Figs. 5D–5G). In addition, FLIM enabled the distinction of nuclear autofluorescence from that of granular autofluorescence. As shown in Fig 3B, young Arabidopsis roots and spinach roots had granular structures with autofluorescence more prominent than that of the nuclei (Figs. 3B), FLIM showed that these granular structures produced significantly longer fluorescence lifetimes (Figs. 5H–5K). Thus, our data demonstrated that live autofluorescence imaging of plant nuclei can be improved using FLIM.

**Figure 5.**
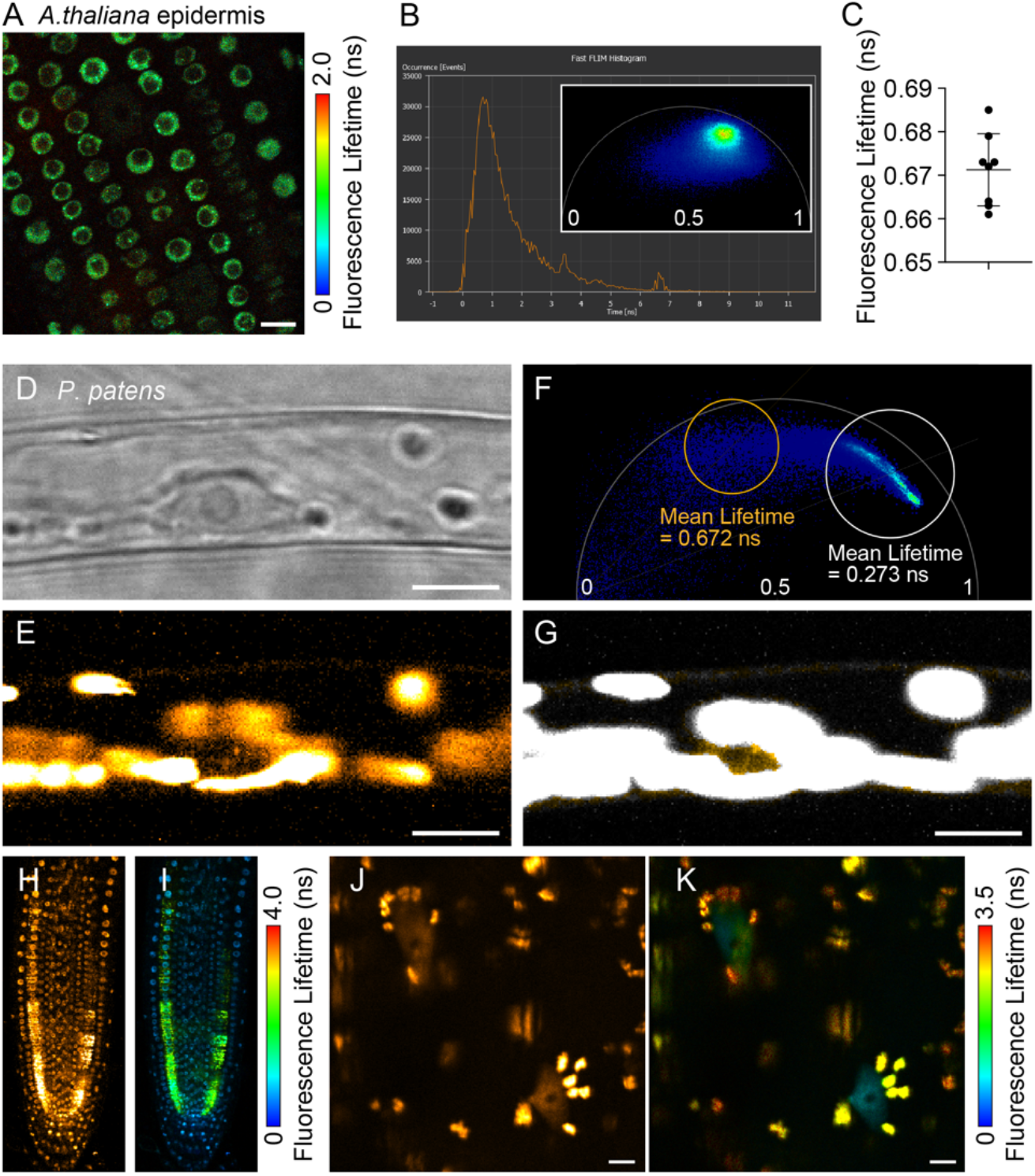
FLIM of the nuclear autofluorescence. (A) FLIM image of *Arabidopsis thaliana* root epidermal cells. (B) A representative histogram of fluorescence lifetime per pixel and a phasor image corresponding to A. (C) Fluorescence lifetime of nuclei. Mean values of fluorescence lifetime of the nuclear NIR autofluorescence in each plant were plotted. Bars represent mean ± SD. *n* = 8 plants. (D, E) Normal confocal images of a *Physcomitrium patens* cell. (D) bright field. (E) autofluorescence excited by 640-nm laser. (F) Phasor plot of fluorescence lifetime in E. (G) FLIM image separated by fluorescence lifetime as indicated in F. Pixels within the orange circle (mean lifetime = 0.672 ns) and white circle (mean lifetime = 0.273 ns) in F were colored in orange and white, respectively. (H) Confocal image of *Arabidopsis thaliana* primary root. (I) FLIM image of H. (J) Confocal image of spinach root. (K) FLIM image of J. *Arabidopsis thaliana* and spinach seedlings were grown on MGRL medium containing 30 µM and 1 µM boric acid, respectively, for 5 days. Scale bars represent 10 µm.

### Phytochromes are responsible for nuclear near-infrared autofluorescence

Since the NIR autofluorescence was predominantly observed in the nucleoplasm, but was likely dispersed during nuclear envelope breakdown (Fig. 1E; Supplementary Fig. S1), we hypothesized that the molecules responsible for this autofluorescence were either intrinsically fluorescent proteins or those conjugated with fluorescent molecules. The nuclear NIR autofluorescence was absent in human HeLa cells and a budding yeast, *Saccharomyces cerevisiae* (Supplementary Figs. S4A–B). In a filamentous fungus, *Aspergillus oryzae*, an intense NIR autofluorescence was observed in the conidia and hyphae, whereas a pattern typical of nuclei was not observed (Supplementary Fig. S4C). Based on our observations, we speculated that the NIR autofluorescence in nucleus is unique to plant species. The phenomenon of fluorescence in plant cells, caused by phytochrome molecules with a peak wavelength of 685 nm, is widely recognized (reviewed in Sineshchekov, 2010). This coincidence has led us to explore the possible involvement of phytochromes in nuclear fluorescence. Given the relatively high expression of *PHYA* and *PHYB* in Arabidopsis (Sharrock and Clack, 2002), we examined the NIR autofluorescence in *phyA/phyB* mutants. Strikingly, the autofluorescence in the nuclei was more reduced in the *phyB* mutant than in the *phyA* mutant (Supplementary Fig. S5), and was completely absent in the *phyA;phyB* double mutant (Figs. 6A and 6B).

**Figure 6.**
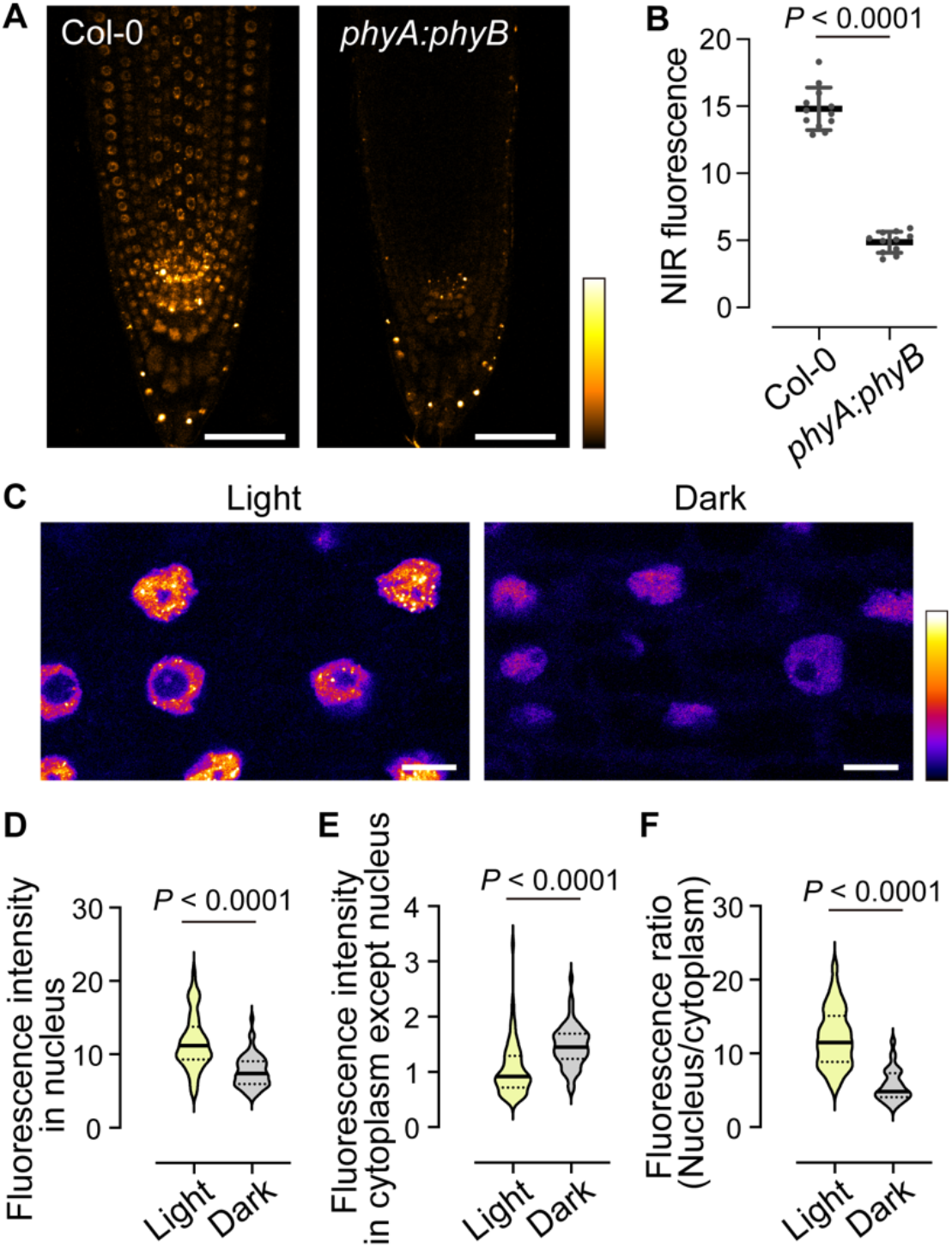
Phytochromes are causative fluorescent proteins for the nuclear NIR fluorescence. (A) NIR autofluorescence excited by 640-nm laser in roots of wild-type Col-0 and *phyA;phyB* double mutant. Plants were grown on modified MGRL medium containing 1 µM boric acid for 4 days. (B) Average NIR autofluorescence intensity from the meristematic to transition zones of the roots (200 µm from the tip) in wild-type and *phyA;phyB*. Horizontal line and error bars represent mean ± SE. *n* = 12 (wild-type) and 11 (*phyA;phyB*) seedlings, respectively. *P* value was calculated by Mann-Whitney test. (C) NIR autofluorescence in root transition zone at 20 h after shifting to light or dark conditions. (D, E) Violin plots representing fluorescence intensity of nucleus (D) and cytoplasm except nucleus (E) in each cell at 20 h after shifting to light and dark conditions. (F) Violin plots representing fluorescence ratio (nucleus/cytoplasm except nucleus) at 20 h after shifting to light and dark conditions. (D–F) *n* = 72 (light) and 60 (dark) cells from >5 different roots, respectively. Solid and broken lines indicate median and quartile lines, respectively. *P* values were calculated by Mann-Whitney test. Scale bars represent 50 µm (A) and 10 µm (C), respectively.

We found that the NIR autofluorescence was observed as foci in the nucleoplasm (Figs. 1E and 1F). This finding is consistent with previous reports that phytochrome undergoes light-dependent transition into so-called photobodies within the nucleoplasm (Van Buskirk et al. 2012; Seluzicki et al. 2017; Pardi and Nusinow, 2021). The subcellular localization of phytochrome is known to influence its function, with nuclear localization decreasing under dark conditions (Salisbury et al. 2007). In agreement with these previous observations, the nuclear NIR autofluorescence was reduced and cytoplasmic NIR autofluorescence was increased in roots acclimated to darkness (Fig. 6C–F). The NIR autofluorescence signals in the nucleus and intranuclear subcompartments were also found to colocalize with GFP-tagged PHYB (Supplementary Fig. S6). Taken together, these results indicate that the nuclear NIR autofluorescence at 656-700 nm arises from phytochrome proteins.

## Discussion

Imaging the nucleus of living cells typically requires labeling the nuclei with either chemical probes or FP-based nuclear markers, with each method possessing its own set of advantages and disadvantages. Chemical labeling necessitates time for the probe to be transported and to stain the target, and chemical probes are eventually degraded, rendering them unsuitable for long-term analysis. DAPI, which is commonly used for staining nuclei in fixed cells, exhibits membrane-impermeability in Arabidopsis cells (Supplementary Fig. S7) and is thus not suitable for live cell imaging. Conversely, FP-based nuclear markers require a process of transient or stable introduction of DNA and sufficient expression of the marker gene in the target cells. One plausible solution for these drawbacks is non-invasive imaging using autofluorescence. Autofluorescence of endogenous biomolecules has been used for live-cell imaging to visualize specific organelles and regions within plant cells. Among the most intense autofluorescence in plants is that derived from chlorophyll. Chlorophylls molecules emit red fluorescence with a bimodal emission, with peaks at 685–695 nm and 720–730 nm, respectively corresponding to photosystem II and photosystem I (Chytilova et al. 2000; Vácha et al. 2007). The autofluorescence of chlorophyll is widely used not only to reveal the localization of chloroplasts but also to measure photosynthetic activity (Baker, 2001; Dutta et al. 2017). Lignin, a phenolic compound in the secondary cell wall, also emits fluorescence, with xylem tissues exhibiting strong green autofluorescence due to their highly-lignified walls (Donaldson, 2020). Additionally, the Casparian strip, a highly-lignified region of cell wall found in mature endodermis of roots, shows a strong autofluorescence, enabling the assessment of its integrity (Naseer et al. 2012). In this study, we present another example of imaging an organelle using autofluorescence without the use of dyes or fluorescent proteins. Nuclear autofluorescence was detected using a conventional confocal microscope, exciting the live sample at 640 nm and detecting emission in the 656–700 nm wavelength range.

We revealed that this NIR autofluorescence was originated from phytochrome, one of the photoreceptors, found in plants, fungi, bacteria, and cyanobacteria. While the NIR fluorescence of plant phytochromes has been previously reported (Sineshchekov, 2010), its suitability for fluorescence imaging had not been evaluated. On the other hand, bacteria phytochromes were also known to exhibit NIR fluorescence and have been utilized to engineer NIR fluorescent proteins for live-cell imaging due to this property (Shu et al. 2009). In recent years, the NIR wavelength range has been used for *in vivo* imaging of plant cells, for instance, a near-infrared fluorescent protein (iRFP) and silicon rhodamine dye (SNAP-Cell 647, SiR647) (Shimizu et al. 2021; Iwatate et al. 2020). Our study cautions that both chloroplast autofluorescence and nuclear autofluorescence must be considered when imaging in this wavelength region.

Near-infrared fluorescence live-imaging has a major disadvantage in imaging in shoots because of the strong autofluorescence derived from chloroplasts. Therefore, the autofluorescence imaging of nuclei is basically limited to roots and pollen grains in which autofluorescence of other organelles is not significant. We found that FLIM enables the distinction between autofluorescence of phytochromes and chlorophylls in *P. patens* (Fig. 5). However, the fluorescence intensity of phytochromes was much weaker than that of chlorophylls, and it is still challenging to image nuclear autofluorescence in Arabidopsis shoots.

The genetic approach successfully validated that both PHYA and PHYB exhibit prominent NIR fluorescence in Arabidopsis cells. In the *phyA* mutant, the intensity of NIR autofluorescence was not considerably different compared to the wild type, whereas it was significantly lower in *phyB*. This observation is consistent with prior results that establish PHYA as a light-labile phytochrome and PHYB as a light-stable phytochrome (Sharrock and Clack, 2002). Interestingly, in Arabidopsis roots germinated in the dark, the NIR autofluorescence of the nucleus was decreased, similar to previous studies, but the NIR fluorescence in the columella region was increased. This enhanced NIR fluorescence was diminished in the *phyA;phyB* double mutants (Supplementary Fig. S8A).

Phytochrome has long been known to be present in the root cap of various crops, such as barley and oat, through immune-antibody staining techniques (Coleman and Pratt, 1974). In maize, *phyA* mRNA accumulates in the columella region in the dark, and its mRNA level decreases in a light-dependent manner (Johnson et al. 1991). In line with previous research, the NIR fluorescence in the columella was absent in Arabidopsis acclimated to light after germination in the dark (Supplementary Fig. S8B). Significantly, NIR autofluorescence imaging revealed the cytoplasmic localization of phytochrome at the subcellular level, which has not been previously demonstrated by immunostaining or *in situ* hybridization methods (Coleman and Pratt, 1974; Johnson et al. 1991). Interestingly, a distinct localization pattern of NIR autofluorescence was observed in certain columella cells, characterized by ring-like structures that likely correspond to the nuclear envelope. This pattern coincided with the localization of the ER-marker, GFP-HDEL. NIR autofluorescence was also observed as punctate structures in the cell periphery (Supplementary Figs. S8C and D). This finding indicates a potential dark-induced sequestration of phytochromes from the nucleus, forming punctate structures on the outside of the nuclear envelope and at the plasma membrane. To elucidate the functional significance of this columella-specific behavior of phytochromes, further investigations are warranted. Our observations also showed in pollen and pollen tubes that NIR autofluorescence was detected solely in the vegetative nucleus (Figs. 2A–C; Video S1). A previous study reported that red light enhances pollen tube elongation in *Arachis hypogaea* (Chhabra and Malik, 1978). The involvement of phytochromes in light-dependent pollen tube elongation in association with the vegetative nucleus can now be easily tested in Arabidopsis using NIR autofluorescence.

Phytochromes are widely conserved in land plants. Nuclear NIR autofluorescence imaging has been demonstrated to be applicable to a wide range of plant species, indicating that investigations into phytochrome protein dynamics in plants are feasible, even in species for which transformation has not been established. Our findings provide a new non-invasive live-cell imaging technique for nuclear dynamics and shed light on the subcellular dynamics of phytochrome, creating a way to expand our understanding of phytochrome biology.

## Experimental Procedures

### Plasmid construction and plant transformation

For the *p35S:nlsGFP* binary vector, a sequence coding the SV40-derived nuclear localization signal PKKKRKVGG (Kalderon et al. 1984) was inserted immediately after the initiation ATG of the GFP sequence by PCR (Table S1). The nlsGFP was inserted into the *Bam*HI and *Kpn*I sites after the *CaMV 35S RNA* promoter (*p35S*) in the pAN19 vector (Miyashima et al. 2011). The resulting construct was digested by *Not*I and integrated into the pBIN40 binary vector by ligation (Miyashima et al. 2011). The *p35S:GFP-TUA5* binary vector was generated using Gateway technology (Invitrogen) by following the Multisite Gateway Technology three-fragment vector construction kit protocol. GFP was inserted into the *Asc*I and *Kpn*I sites after the *p35S* sequence in the p35S-p4p1r entry vector. The genomic region from the start to stop codon of TUA5 (AT5G19780) was cloned into pDONR221. Three entry clones, p35S-EGFP-p4p1r, TUA5- pDONR221 and t35S-p2rp3 were transferred into the binary vector pBm43GW (Karimi et al. 2005). The entry vectors were kindly provided by Dr. A. Untergasser (University of Heidelberg, Germany). The final binary vectors were introduced into *Agrobacterium tumefaciens* strain GV3101:pMP90 and used to carry out the transformation of Arabidopsis.

### Plant materials

Arabidopsis transgenic line expressing *mCherry-NIP5;1genomic* and *BOR1-EGFP* (Takano et al., 2017) was kindly provide by Dr. Junpei Takano (Osaka Metropolitan University, Japan). Arabidopsis plants expressing *H2B-mClover* (Kurihara et al., 2015) and *GFP-HDEL* (Mitsuhashi et al., 2000) were kindly provided by Dr. Daisuke Kurihara and Dr. Yoko Mizuta (Nagoya University, Japan). Seeds of *Arabidopsis thaliana* Col-0 ecotype and *Nicotiana benthamiana* were our laboratory stock. Seeds of *Oryza sativa* subsp. *japonica* ‘Nipponbare’ were kindly provided by Dr. Motoyuki Ashikari (Nagoya University, Japan). For colocalization analysis between NIR autofluorescence and PHYB-GFP, transgenic lines expressing PHYB-GFP under the control of either its own promoter or the cauliflower mosaic virus 35S RNA promoter were grown from seeds of *proPHYB:PHYB-GFP* in *phyB-1* (Bpro7) (Endo et al. 2005) or *pro35S:PHYB-GFP* in *phyB-5* (PBG3-1) (Matsushita et al. 2003) provided by Dr. Tomonao Matsushita (Kyoto University, Japan). Seeds of *Capsella bursa-pastoris* and *Cardamine hirsuta* were collected from plants grown naturally in Nagoya, Aichi, Japan. Seeds of *Eruca vesicaria*, *Daucus carota*, *Coriandrum sativum*, *Spinacia oleracea*, and *Cucumis sativus* marketed for home gardens and *Allium cepa* were purchased in a local grocery store. Plants were grown on either half-strength Murashige Skoog (MS) medium (Murashige and Skoog, 1962) supplemented with 1% (w/v) sucrose and 1% (w/v) agar or modified MGRL medium (Yoshinari and Takano, 2020) containing 0.5, 1 or 30 µM boric acid, 1% (w/v) sucrose, and 1.5% (w/v) gellan gum. Seedlings were grown in a growth chamber equipped with a broad-range LED light (NK System) set to either a 16-/8-h light/dark cycle at 23°C, for *Arabidopsis thaliana*, *Eruca vesicaria*, *Capsella bursa-pastoris* and *Cardamine hirsuta,* or 12-/12-h light/dark cycle at 25°C for *Daucus carota*, *Coriandrum sativum*, *Spinacia oleracea*, and *Cucumis sativus*. For the analysis of NIR autofluorescence in dark-grown seedlings, medium-containing plates were incubated under a light condition in the growth chamber for 24 hours and then covered with aluminum foil. Tobacco BY-2 cells were cultured in modified Linsmaier and Skoog (LS) medium (Katsuta et al. 1990) buffered with 0.05% (w/v) 2-(N-morpholino)ethanesulfonic acid (MES) at pH 5.8. Cells were subcultured once a week by adding 1.5 ml of cell suspension into 80 ml of fresh medium. Cultures were maintained in the dark at 25°C with shaking (120 rpm). The moss *Physcomitrium patens* strain Cove-NIBB strain was cultured 7-8 days on BCDAT medium under continuous white light at 25℃ (Nishiyama et al, 2000).

### HeLa cell cultures

HeLa cells were grown in Dulbecco’s Modified Eagle’s medium (D-MEM) containing L-glutamate, D-glucose, phenol red (Fujifilm Wako) and supplemented with 10% (v/v) fetal bovine serum (FB- 1061/500, Biosera), and 1% antibiotics (penicillin/streptomycin) (Fujifilm Wako) in a humidified incubator at 37°C under 5% CO2 atmosphere. HeLa cells were passaged in D-MEM containing L-glutamate and D-glucose without phenol red (Fujifilm Wako) 2 days before live-cell imaging.

### Fungal materials

*Saccharomyces cerevisiae* BY4741 strain and *Aspergillus oryzae* were grown on YPD media containing 2% agar in an incubator at 30°C. *Aspergillus oryzae* was a product prepared for food fermentation and was purchased at a local grocery store in Japan.

### Pollen grains and germination

Pollen grains were collected from Col-0 plants and germinated as described previously (Isoda et al. 2022). For pollen nucleus staining, pollen grains were incubated either with Cellstain® Hoechst33342 solution (Cat#: H342, Dojindo) diluted to 1 µg/mL with Milli-Q ultra-pure water for 5 min at room temperature or with a pollen germination medium containing 2000-fold SYBR Green I (Cat#: 5760A, Takara) and 5 µg/mL DAPI (Cat# 10236276001, Roche).

### DAPI staining

To stain the nuclei of seedlings, four-day-old whole seedlings were submerged in PBS buffer containing 4% (w/v) paraformaldehyde at room temperature for 1 h to fix the cells. The specimens were washed four times with PBS buffer, stained with 2 µg/mL DAPI diluted in PBS for 20 min, and washed once with PBS buffer. For the staining of cell walls in living cells, four-day-old seedlings were submerged in MGRL liquid medium containing 2 µg/mL DAPI for 30 min and washed once with MGRL liquid medium.

### Confocal laser scanning microscopy

For fluorescence imaging, a Zeiss LSM800 equipped with 20× dry (0.8 NA Plan-Apochromat, Zeiss), 40× water-immersion (1.1 NA LD C-Apochromat, Zeiss), and 63× oil-immersion (1.4 NA Plan-Apochromat, Zeiss) objective lenses and 405-, 488-, 561-, and 640-nm lasers was used. For nuclear autofluorescence imaging, the excitation wavelength was set at 640 nm and the detection was set at 656-700 nm. For the time-gating, lamda (λ)-scanning, and fluorescence lifetime microscopy (FLIM) analyses, a Leica TCS-SP8 gSTED equipped with 20× (0.75 NA HC PL APO CS2, Leica) and 63× oil-immersion (1.40 NA HC PL APO CS2, Leica) objectives and a white-light laser was used. For λ-scanning, autofluorescence was excited by a laser at 640 nm, and the fluorescence was scanned with a 10-nm detection window from 645 nm to 795 nm without any gaps (15 steps). For time-lapse imaging, plants were placed on a glass-bottom dish and covered by MGRL solid medium used for the plant culture. Time-lapse imaging of pollen germination was performed using a Zeiss LSM800, with images taken at 5-min intervals using a Z-stack in 1.2-µm increments, employing the 63× oil-immersion objective lens. For fluorescence lifetime imaging, a Leica SP8-FALCON equipped with an HC PL APO CS2 63×/1.40 objective lens and a white-light laser was used. The nuclear autofluorescence was excited at 640 nm and collected between 650– 670 nm. The phasor plot was used for detecting the fluorescence lifetime population of nuclear autofluorescence by LAS X software (Uno et al. 2021).

### Statistics

All statistical analyses were done by Prism 8 for macOS ver. 8.4.3 (GraphPad Software). Statistical methods were indicated in each figure legend.

## Data Availability

The data underlying this article are available in the article.

## Accession numbers

AT2G47160 (BOR1), AT4G10380 (NIP5;1), AT5G19780 (TUA5), AT1G09570 (PHYA), AT2G18790 (PHYB)

## Acknowledgements

We would like to thank Toshinori Kinoshita (Nagoya University) and Winslow Briggs (Carnegie Institution for Science) for providing *phyA* (*phyA-211*), *phyB* (*phyB-9*), and *phyA;phyB* seeds; Tomonao Matsushita (Kyoto University) for providing Bpro7 and PBG3-1 seeds; Daisuke Kurihara (ITbM, Nagoya University) for providing H2B-mClover seeds; and Junpei Takano (Osaka Metropolitan University) for providing BOR1-GFP mCherry-NIP5;1 seeds. We are grateful to Keisuke Nagai and Motoyuki Ashikari (Nagoya University) for providing Nipponbare seeds. We also thank Kayoko Hirano (Nagoya University) for excellent technical assistance. This work was supported by Advanced Bioimaging Support and the Program for Promoting the Enhancement of Research Universities as young researcher units for the advancement of new and undeveloped fields at Nagoya University.

## Author Contributions

A.Y. and M.N. conceived the project. A.Y., R.I., N.Y., Y.S., and M.N. performed experiments. J.J.L. and D.W.E. contributed materials. A.Y., R.I., N.Y., Y.S., W.B.F., and M.N. analyzed data, and A.Y., R.I., Y.S., W.B.F., and M.N. wrote the paper.

## Short legends for Supporting Information

## Reference

Baker, N.R. (2001). High resolution imaging of photosynthetic activities of tissues, cells and chloroplasts in leaves. Journal of Experimental Botany 52: 615–621.

Chhabra, N. and Malik, C.P. (1978). Influence of Spectral Quality of Light on Pollen Tube Elongation in Arachis hypogaea. Annals of Botany 42: 1109–1117.

Chytilova, E., Macas, J., Sliwinska, E., Rafelski, S.M., Lambert, G.M., and Galbraith, D.W. (2000). Nuclear Dynamics in Arabidopsis thaliana. MBoC 11: 2733–2741.

Coleman, R.A. and Pratt, L.H. (1974). Subcellular localization of the red-absorbing form of phytochrome by immunocytochemistry. Planta 121: 119–131.

Donaldson, L. (2020). Autofluorescence in Plants. Molecules 25: 2393.

Dundr, M. and Misteli, T. (2001). Functional architecture in the cell nucleus. Biochemical Journal 356: 297–310.

Dutta, S., Cruz, J.A., Imran, S.M., Chen, J., Kramer, D.M., and Osteryoung, K.W. (2017). Variations in chloroplast movement and chlorophyll fluorescence among chloroplast division mutants under light stress. Journal of Experimental Botany 68: 3541–3555.

Endo, M., Nakamura, S., Araki, T., Mochizuki, N., and Nagatani, A. (2005). Phytochrome B in the Mesophyll Delays Flowering by Suppressing *FLOWERING LOCUS T* Expression in Arabidopsis Vascular Bundles. Plant Cell 17: 1941–1952.

Isoda, R., Palmai, Z., Yoshinari, A., Chen, L.-Q., Tama, F., Frommer, W.B., and Nakamura, M. (2022). SWEET13 transport of sucrose, but not gibberellin, restores male fertility in *Arabidopsis sweet13;14*. Proc. Natl. Acad. Sci. U.S.A. 119: e2207558119.

Iwatate, R.J., Yoshinari, A., Yagi, N., Grzybowski, M., Ogasawara, H., Kamiya, M., Komatsu, T., Taki, M., Yamaguchi, S., Frommer, W.B., and Nakamura, M. (2020). Covalent Self-Labeling of Tagged Proteins with Chemical Fluorescent Dyes in BY-2 Cells and Arabidopsis Seedlings. Plant Cell 32: 3081–3094.

Johnson, E.M., Pao, L.I., and Feldman, L.J. (1991). Regulation of Phytochrome Message Abundance in Root Caps of Maize: Spatial, Environmental, and Genetic Specificity. Plant Physiol. 95: 544–550.

Kalderon, D., Richardson, W.D., Markham, A.F., and Smith, A.E. (1984). Sequence requirements for nuclear location of simian virus 40 large-T antigen. Nature 311: 33–38.

Kapuscinski, J. (1995). DAPI: a DNA-Specific Fluorescent Probe. Biotechnic & Histochemistry 70: 220–233.

Karimi, M., De Meyer, B., and Hilson, P. (2005). Modular cloning in plant cells. Trends in Plant Science 10: 103–105.

Katsuta, J., Hashiguchi, Y., and Shibaoka, H. (1990). The role of the cytoskeleton in positioning of the nucleus in premitotic tobacco BY-2 cells. Journal of Cell Science 95: 413–422.

Ketelaar, T., Faivre-Moskalenko, C., Esseling, J.J., De Ruijter, N.C.A., Grierson, C.S., Dogterom, M., and Emons, A.M.C. (2002). Positioning of Nuclei in Arabidopsis Root Hairs: An Actin-Regulated Process of Tip Growth. Plant Cell 14: 2941–2955.

Kurihara, D., Mizuta, Y., Sato, Y., and Higashiyama, T. (2015). ClearSee: a rapid optical clearing reagent for whole-plant fluorescence imaging. Development: dev.127613.

Latt, S.A. and Wohlleb, J.C. (1975). Optical studies of the interaction of 33258 Hoechst with DNA, chromatin, and metaphase chromosomes. Chromosoma 52: 297–316.

Matsushita, T., Mochizuki, N., and Nagatani, A. (2003). Dimers of the N-terminal domain of phytochrome B are functional in the nucleus. Nature 424: 571–574.

Mitsuhashi, N., Shimada, T., Mano, S., Nishimura, M., and Hara-Nishimura, I. (2000). Characterization of Organelles in the Vacuolar-Sorting Pathway by Visualization with GFP in Tobacco BY-2 Cells. Plant and Cell Physiology 41: 993–1001.

Miyashima, S., Koi, S., Hashimoto, T., and Nakajima, K. (2011). Non-cell-autonomous microRNA165 acts in a dose-dependent manner to regulate multiple differentiation status in the *Arabidopsis* root. Development 138: 2303–2313.

Murashige, T. and Skoog, F. (1962). A Revised Medium for Rapid Growth and Bio Assays with Tobacco Tissue Cultures. Physiol Plant 15: 473–497.

Naseer, S., Lee, Y., Lapierre, C., Franke, R., Nawrath, C., and Geldner, N. (2012). Casparian strip diffusion barrier in *Arabidopsis* is made of a lignin polymer without suberin. Proc. Natl. Acad. Sci. U.S.A. 109: 10101–10106.

Pardi, S.A. and Nusinow, D.A. (2021). Out of the Dark and Into the Light: A New View of Phytochrome Photobodies. Front. Plant Sci. 12: 732947.

Pronina, T., Pavlova, E., Dil’mukhametova, L., and Ugrumov, M. (2022). Development of the Periventricular Nucleus as a Brain Center, Containing Dopaminergic Neurons and Neurons Expressing Individual Enzymes of Dopamine Synthesis. IJMS 23: 14682.

Salisbury, F.J., Hall, A., Grierson, C.S., and Halliday, K.J. (2007). Phytochrome coordinates Arabidopsis shoot and root development. Plant J 50: 429–438.

Seluzicki, A., Burko, Y., and Chory, J. (2017). Dancing in the dark: darkness as a signal in plants: Dark signaling in plants. Plant, Cell & Environment 40: 2487–2501.

Sharrock, R.A. and Clack, T. (2002). Patterns of Expression and Normalized Levels of the Five Arabidopsis Phytochromes. Plant Physiology 130: 442–456.

Shimizu, Y. et al. (2021). Cargo sorting zones in the trans-Golgi network visualized by super-resolution confocal live imaging microscopy in plants. Nat Commun 12: 1901.

Shu, X., Royant, A., Lin, M.Z., Aguilera, T.A., Lev-Ram, V., Steinbach, P.A., and Tsien, R.Y. (2009). Mammalian Expression of Infrared Fluorescent Proteins Engineered from a Bacterial Phytochrome. Science 324: 804–807.

Sineshchekov, V.A. (2010). Fluorescence and Photochemical Investigations of Phytochrome in Higher Plants. Journal of Botany 2010: 1–15.

Takano, J., Yoshinari, A., and Luu, D.-T. (2017). Plant Aquaporin Trafficking. In Plant Aquaporins, F. Chaumont and S.D. Tyerman, eds, Signaling and Communication in Plants. (Springer International Publishing: Cham), pp. 47–81.

Tamura, K., Iwabuchi, K., Fukao, Y., Kondo, M., Okamoto, K., Ueda, H., Nishimura, M., and Hara-Nishimura, I. (2013). Myosin XI-i Links the Nuclear Membrane to the Cytoskeleton to Control Nuclear Movement and Shape in Arabidopsis. Current Biology 23: 1776–1781.

Uno, K., Sugimoto, N., and Sato, Y. (2021). N-aryl pyrido cyanine derivatives are nuclear and organelle DNA markers for two-photon and super-resolution imaging. Nat Commun 12: 1–9.

Vácha, F., Sarafis, V., Benediktyová, Z., Bumba, L., Valenta, J., Vácha, M., Sheue, Ch.-R., and Nedbal, L. (2007). Identification of Photosystem I and Photosystem II enriched regions of thylakoid membrane by optical microimaging of cryo-fluorescence emission spectra and of variable fluorescence. Micron 38: 170–175.

Van Buskirk, E.K., Decker, P.V., and Chen, M. (2012). Photobodies in Light Signaling. Plant Physiology 158: 52–60.

von Wangenheim, D., Hauschild, R., Fendrych, M., Barone, V., Benková, E., and Friml, J. Live tracking of moving samples in confocal microscopy for vertically grown roots. eLife 6: e26792.

Yagi, N., Yoshinari, A., Iwatate, R.J., Isoda, R., Frommer, W.B., and Nakamura, M. (2021). Advances in Synthetic Fluorescent Probe Labeling for Live-Cell Imaging in Plants. Plant Cell Physiol 62: 1259–1268.

Yoshinari, A. and Takano, J. (2020). Analysis of and Intracellular Trafficking of Boric Acid/Borate Transport Proteins in Arabidopsis. In Plant Endosomes, M.S. Otegui, ed, Methods in Molecular Biology. (Springer US: New York, NY), pp. 1–13.

Yu, H.-G., Hiatt, E.N., Chan, A., Sweeney, M., and Dawe, R.K. (1997). Neocentromere- mediated Chromosome Movement in Maize. J Cell Biol 139: 831–840.

