## Supplementary Information for "Near-infrared imaging of phytochrome-derived autofluorescence in plant nuclei"

**This PDF file includes:**

Figures S1 to S8
Tables S1
Legends of Supporing Videos

### Supplementary Figure S1

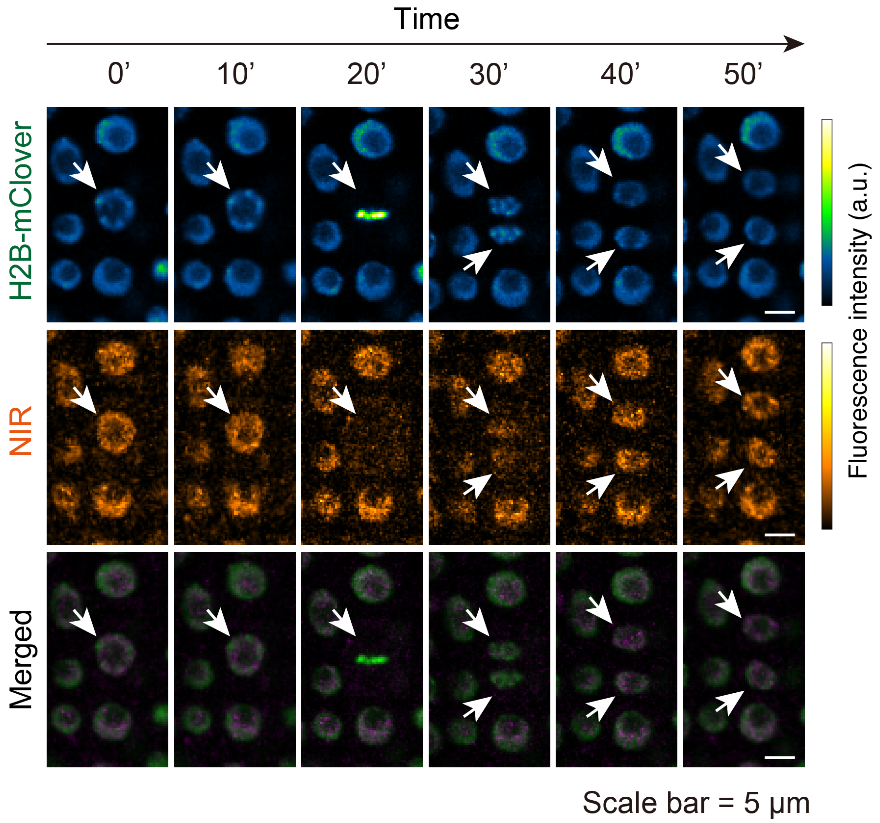


**Figure S1. Near-infrared autofluorescence imaging of dividing cell**

Time-lapse confocal imaging of epidermal cells in root meristematic zone of transgenic *Arabidopsis thaliana* expressing H2B-mClover. Near-infrared autofluorescence was excited by 640-nm laser. Arrows indicate mitotic cell and daughter cells. Plant was grown on modified MGRL medium containing 30 μM boric acid for 3 days. Scale bars represent 5 μm.

**Supplementary Figure S2**

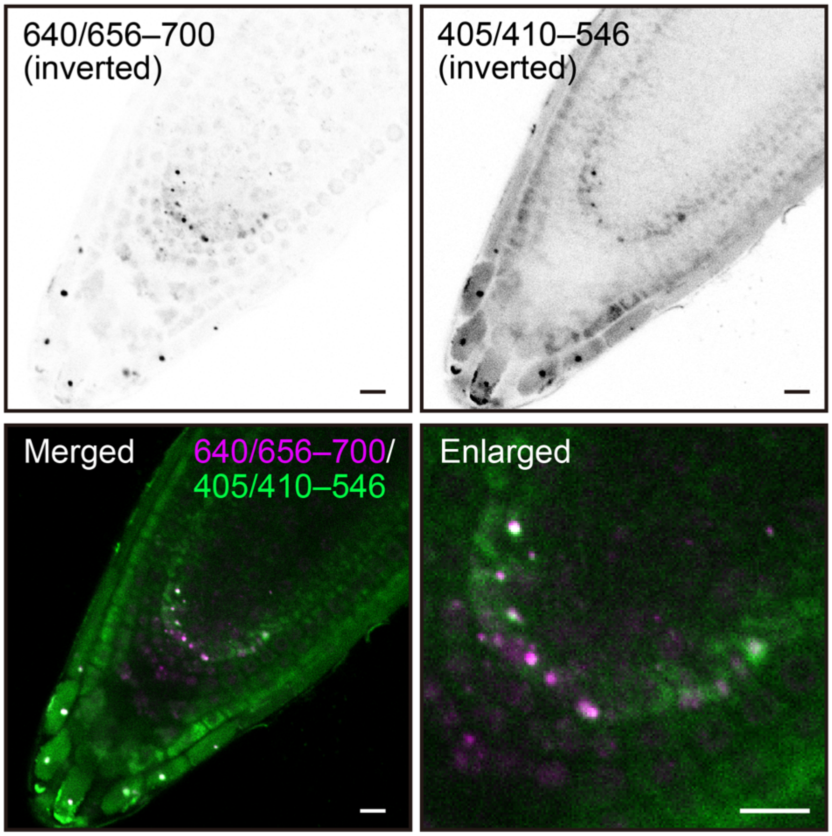


**Figure S2. Root tip emits distinct patterns of autofluorescence**

Inverted confocal images of two distinct autofluorescence in wild-type Col-0 plant. Fluorescence was taken by different excitation/detection combinations, 640/656–700 and 405/410–546. Punctate structures smaller than nucleus found in each image condition were well overlapped. Scale bars represent 10 μm. Plant was grown on half-strength of MS medium for 4 days. Scale bars represent 100 μm.

### Supplementary Figure S3

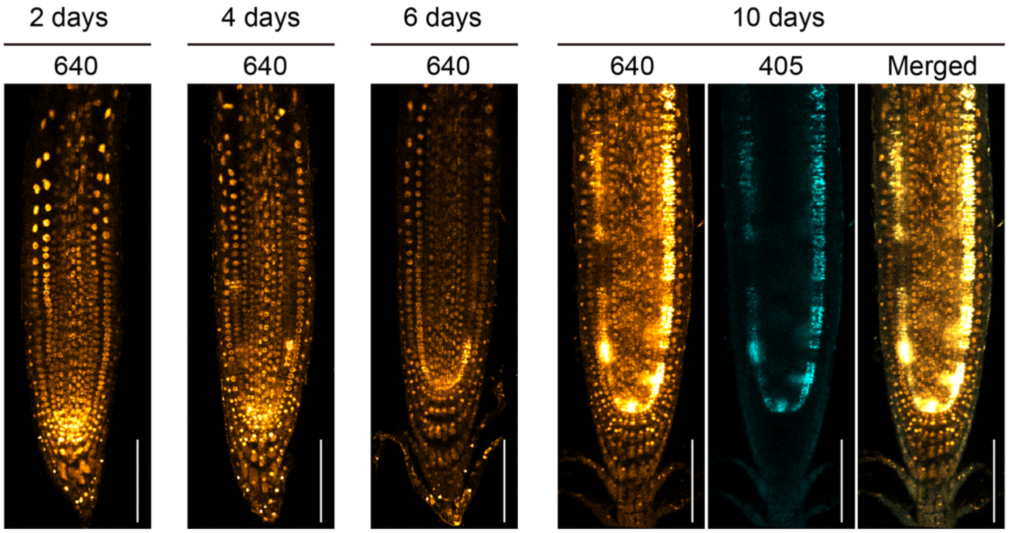


**Figure S3. Old plant roots emit a strong autofluorescence in the cortex**

Confocal images of autofluorescence excited by 640 or 405 nm in wild-type Col-0 plants. Number above the images indicate the excitation wavelength. Ten-day-old root showed a strong NIR fluorescence in the cortex, which was also observed at 405 nm excitation. Scale bars represent 50 µm.

### Supplementary Figure S4

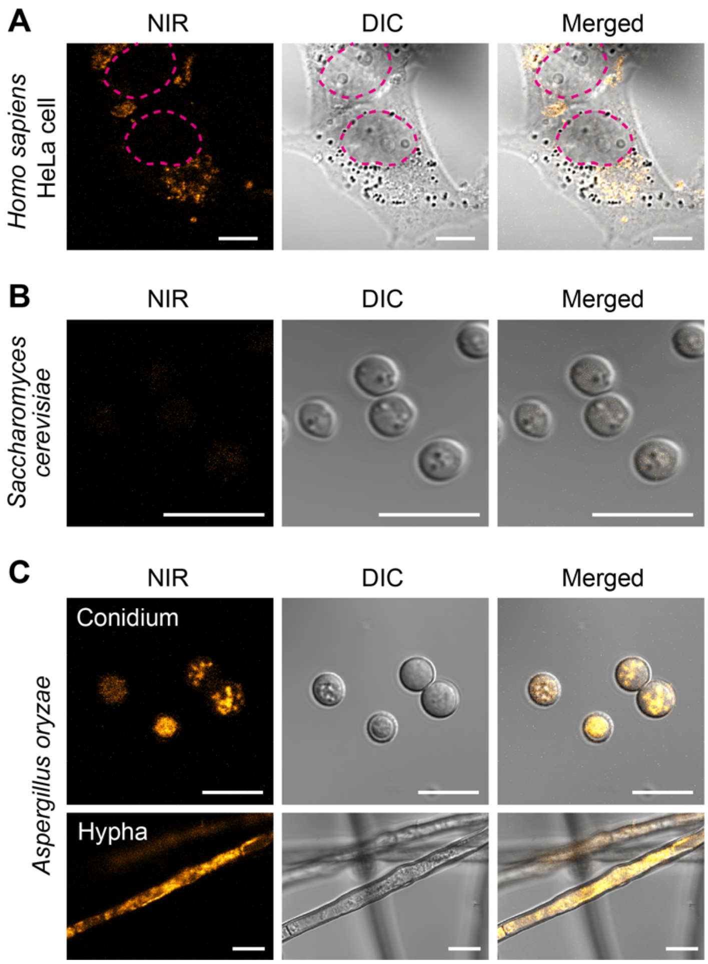


**Figure S4. Near-infrared autofluorescence in mammal and fungal cells**

Confocal images of NIR fluorescence excited by 640-nm laser in *Homo sapiens* HeLa cells, *Saccharomyces cerevisiae* BY4741, and *Aspergillus oryzae*. (A) HeLa cells cultured in D-MEM medium without phenol red that potentially causes optical crosstalk with the NIR fluorescence. Circles indicate nuclei. (B) Yeast cells. (C) Conidia and Hyphae of *Aspergillus oryzae*. Scale bars represent 10 μm.

### Supplementary Figure S5

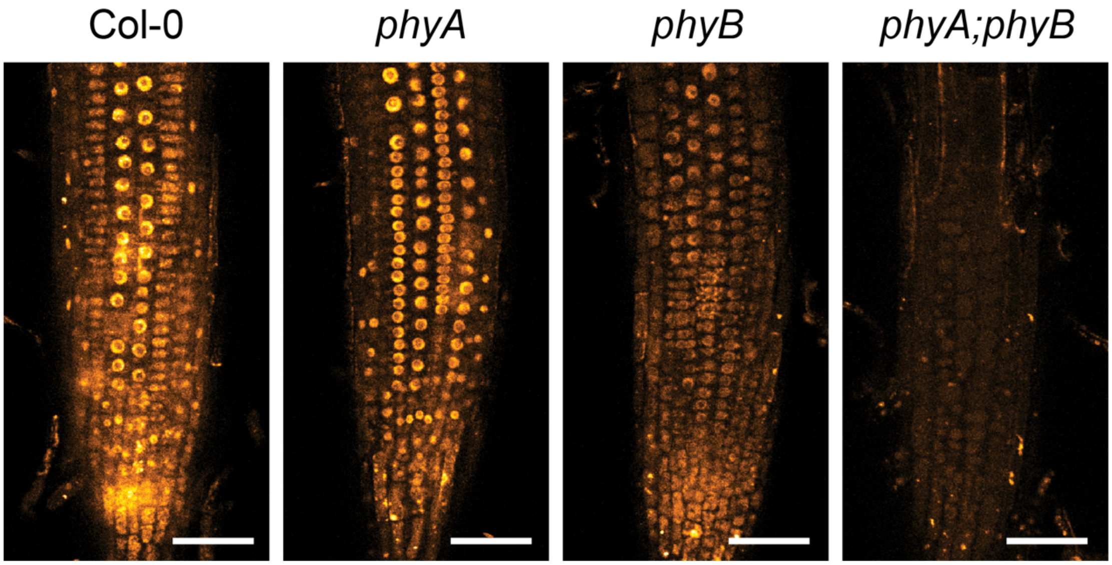


**Figure S5. Near-infrared autofluorescence in mutants lacking phytochromes**

Confocal images of nuclear NIR fluorescence excited by 640-nm laser in wild-type Col-0, *phyA*, *phyB*, and *phyA;phyB* double mutants. Plants were grown on half-strength of MS medium for 4 days. Scale bars represent 50 µm.

**Supplementary Figure S6**

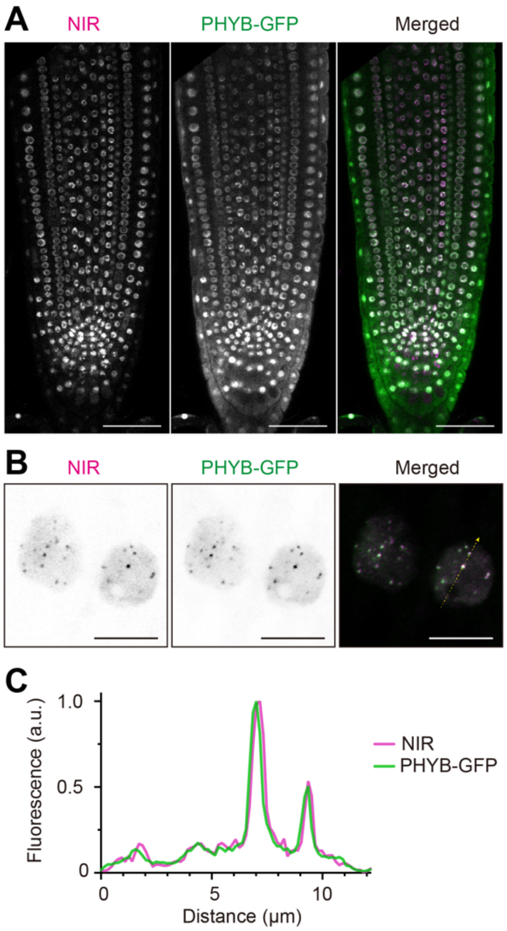


**Figure S6. Nuclear autofluorescence overlaps with PHYB-GFP fluorescence**

(A) Confocal image of NIR and PHYB-GFP fluorescence in the root of 4-day-old *ProPHYB:PHYB-GFP/phyB-1* transgenic plant (Bpro7). (B) Confocal image of NIR and PHYB-GFP fluorescence in the root of 4-day-old *Pro35S:PHYB-GFP/phyB-5* transgenic plant (PBG3-1). (C) Fluorescence intensity of NIR and PHYB-GFP along the arrow indicated in (B). Plants were treated with a normal light condition. Scale bars represent 50 μm (A) and 10 μm (B), respectively.

### Supplementary Figure S7

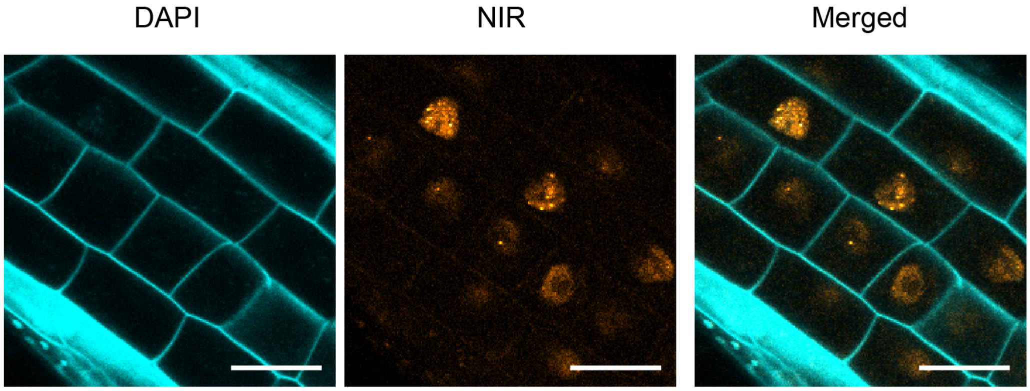


**Figure S7. DAPI staining of living cells in root**

Confocal images of epidermal cells in transition zone of *Arabidopsis thaliana* root. Plant was grown on modified MGRL medium containing 0.5 μM boric acid for 4 days and stained with 2 μg/mL DAPI for 30 min.

### Supplementary Figure S7

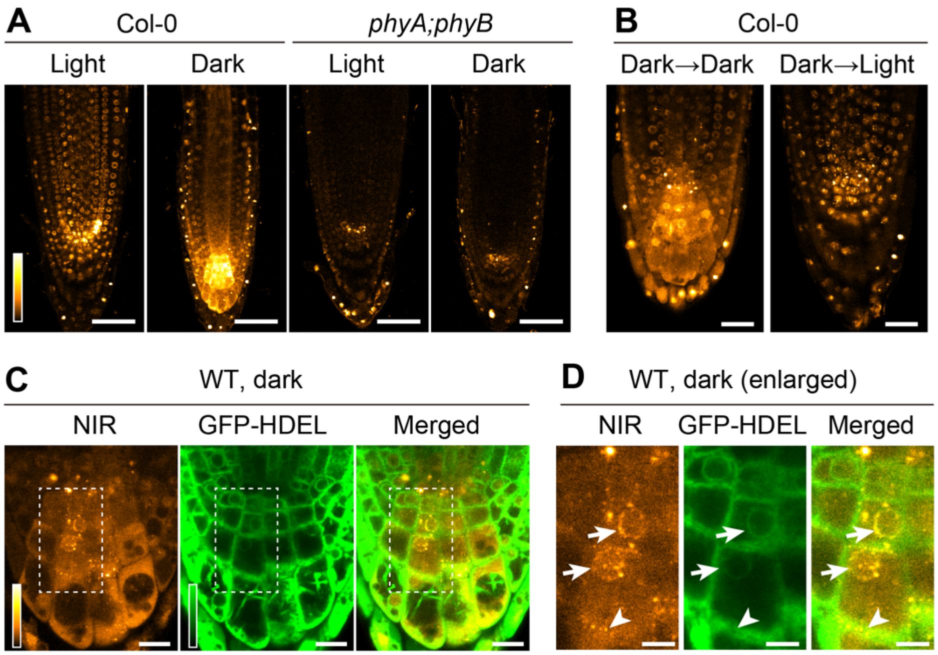


**Figure S8. Near-infrared autofluorescence in plants grown under dark condition**

(A) Near-infrared autofluorescence excited by 640-nm laser in root tip of wild-type Col-0 and *phyA;phyB* mutants grown under light and dark conditions. (B) NIR fluorescence in dark-grown seedling after shifting to a dark or light condition for 270 min. (C) Confocal images of transgenic *Arabidopsis thaliana* expressing an ER marker, GFP-HDEL. Near-infrared autofluorescence excited by 640-nm laser. (D) Enlarged images of the indicated area in C. Arrows and arrowhead indicate fluorescence observed in ring-like structures which overlaps with GFP-HDEL fluorescence and in cell periphery, respectively. Scale bars represent 50 μm (A), 20 μm (B), 10 μm (C), and 5 μm (D), respectively.

**Table S1.** Primers used in the study

| Name | Sequence (5' -> 3') | Purpose |
|---|---|---|
| BamHI_nlsGFP_F | GGATCCATGCTGCAGCCTAAGAAGAAGAGAAAGGTTGG | Add NLS sequence and cloning into pAN19 |
| GFP_KpnI_R | GGTACCTCACTTGTACAGCTCGTCCATG | |
| AcsI_EGFP_F | GGCGCGCCATGGTGAGCAAGGGCGAGGAGCTG | Subcloning of EGFP sequence |
| EGFP_KpnI_R | TGGTACCCTTGTACAGCTCGTCCATGCCGAG | |

**Legends of Supporting Videos**

**Video S1. Time-lapse imaging of nuclear dynamics in root hair elongation of Arabidopsis.**
Movie of time-lapse confocal images of elongating root hair of transgenic *Arabidopsis thaliana* plant expressing H2B-mClover. Increment between frames was 5 min. Scale bar represents 50 µm.

**Video S2. Time-lapse imaging of nuclear dynamics in pollen tube elongation.**
Movie of time-lapse confocal images of elongating pollen tubes. Pollen grains of wild-type Col-0 plant were placed on pollen germination medium containing SYBR Green I and incubated. SYBR Green I (left), NIR autofluorescence (center), and merged images with DIC (right). Scale bar represents 10 µm.

**Video S3. Time-lapse imaging of root elongation of *Cardamine hirsute*.**
Time-lapse imaging of NIR autofluorescence in 4-day-old *Cardamine hirsute* root. Scale bar represents 100 µm.

**Video S4. Time-lapse imaging of nuclear dynamics in Roquette root.**
Time-lapse imaging of NIR autofluorescence in transition zone of Roquette (*Eruca vesicaria* subsp. *Sativa*) root. Scale bar represents 50 µm.

**Video S5. Time-lapse imaging of nuclear migration in Roquette root hair.**
Time-lapse imaging of NIR autofluorescence in root hair of Roquette (*Eruca vesicaria* subsp. *Sativa*). Scale bar represents 50 µm.
